# Pilot Study of Metabolomic Clusters as State Markers of Major Depression and Outcomes to CBT Treatment

**DOI:** 10.1101/658906

**Authors:** Sudeepa Bhattacharyya, Boadie W. Dunlop, Siamak Mahmoudiandehkordi, Ahmed T. Ahmed, Gregory Louie, Mark A. Frye, Richard M. Weinshilboum, Ranga R Krishnan, A John Rush, Helen S. Mayberg, W. Edward Craighead, Rima Kaddurah-Daouk

## Abstract

Major depressive disorder (MDD) is a common and disabling syndrome with multiple etiologies that is defined by clinically elicited signs and symptoms. In hopes of developing a list of candidate biological measures that reflect and relate closely to the severity of depressive symptoms, so-called “state-dependent” biomarkers of depression, this pilot study explored the biochemical underpinnings of treatment response to cognitive behavior therapy (CBT) in medication-freeMDD outpatients. Plasma samples were collected at baseline and week 12 from a subset of MDD patients (N=26) who completed a course of CBT treatment as part of the Predictors of Remission in Depression to Individual and Combined Treatments (PReDICT) study. Targeted metabolomic profiling using the the AbsoluteIDQ^®^ p180 Kit and LC-MS identified eight “co-expressed” metabolomic modules. Of these eight, three were significantly associated with change in depressive symptoms over the course of the 12-weeks. Metabolites found to be most strongly correlated with change in depressive symptoms were branched chain amino acids, acylcarnitines, methionine sulfoxide, and α-aminoadipic acid (negative correlations with symptom change) as well as several lipids, particularly the phosphatidlylcholines (positive correlation). These results implicate disturbed bioenergetics as an important state marker in the pathobiology of MDD. Exploratory analyses contrasting remitters to CBT versus those who failed the treatment further suggest these metabolites may serve as mediators of recovery during CBT treatment. Larger studies examining metabolomic change patterns in patients treated with pharmacotherapy or psychotherapy will be necessary to elucidate the biological underpinnings of MDD and the -specific biologies of treatment response.

## 1. INTRODUCTION

Major depressive disorder (MDD) is a clinical syndrome that has multiple etiologies, and responds to a diverse range of treatments that affect various biological pathways. Nevertheless, it is highly likely that specific biological processes underpin the clinical presentation of the disorder. Identifying these *state* biological processes could provide a more precise gauge of the pathophysiological processes underpinning the clinical-symptomatic expression of MDD, and that also could reflect treatments’ biological effect beyond the information gleaned from the simple assessment of signs and symptoms (Rush and Ibrahim 2018)

Biological, physiological, neuro-functional, and other measures that are most closely tied to and reflective of the *clinical expression* of MDD or to the symptomatic expression of other medical syndromes are often referred to “state-dependent” markers (Rush and Ibrahim, 2018). Central venous pressure (CVP), for example, is a state-dependent measure for congestive heart failure (CHF). The greater the CVP, the more severe the symptoms that define CHF such as pedal edema, pulmonary effusion, orthopnea, and dyspnea. On the other hand, “trait-like” markers, are those measures that are persistently abnormal both during and between clinically symptomatic episodes (Rush and Ibrahim 2018). “Trait-like” markers often reflect the underlying pathobiology of the condition that either sets the stage for the initial clinical expression of the disorder or that reflect the effect/consequence of the clinical episode itself even after the episode ends. The latter are sometimes said to be “scars” or consequences of the clinical episode. Left or right ventricular hypertrophy, for instance, can be consequences of repeated episodes of congestive heart failure (Senni and Redfield 1997). For MDD, hypercortisolemia is known to be highly state dependent in psychotic or melancholic depressions (Pariante 2017). On the other hand, some sleep EEG parameters appear to be more trait-like (i.e. persistent even between clinically apparent major depressive episodes) than state-dependent (only apparent during clinically apparent depressive episodes) (Thase et al. 1998; Kraemer et al. 1994). However, neither state nor trait markers for MDD have been found that are as yet of sufficient value to enter clinical practice.

Metabolomics have the potential to define specific biochemical processes that underpin MDD, the effect of treatment on the biological processes that underpin MDD, and the effects of treatments on the biology of MDD. To date, however, metabolomic profiling has largely been conducted with depressed patients who have been taking antidepressant medications that are known to affect metabolomics profiles (Abo et al. 2012; Zhu et al. 2013; Kaddurah-Daouk et al. 2013; Kaddurah-Daouk et al. 2011; Rotroff et al. 2016); these medications directly interfere with the identification of state dependent measures.

Pharmacometabolomic studies from our group have previously reported perturbations in intermediates of TCA cycle, urea cycle, amino acids, and lipids in depressed patients exposed to sertraline (Zhu et al. 2013; Kaddurah-Daouk et al. 2013; Kaddurah-Daouk et al. 2011). Another study utilizing intravenous ketamine treatment in depressed patients reported changes in tryptophan metabolism, acylcarnitines, urea cycle intermediates, and lipid metabolism (Rotroff et al. 2016). In a cross-sectional study, the branched chain amino acids (BCAAs) Valine, Leucine, and Isoleucine were significantly lower in MDD patients compared to healthy controls and were negatively correlated to Hamilton Depression Rating Scale scores. In a rat model of depression, biogenic amines like putrescine, spermine, and spermidine were significantly reduced in the hippocampus of stressed animals compared to non-stressed ones, but the biogenic amines were restored by the antidepressant effect of S-adenosyl-L-methionine (Genedani et al. 2001). Plasma lipid and acylcarnitine profiles, which have also been implicated in animal models of depression, suggest inflammatory conditions and incomplete mitochondrial β-oxidations as primary phenomena associated with the pathophysiology of MDD(Chen et al. 2014).

This pilot study utilized a sample from the cognitive behavior therapy (CBT) arm of theEmory Predictors of Remission in Depression to Individual and Combined Treatments (PReDICT) study, a randomized controlled trial of previously untreated patients with MDD (Dunlop, Kelley, et al. 2017). This sample avoids the likely confounding effects of medications on endogenous metabolomic processes under study; i.e., the sample ensures that when patients improve symptomatically- (or not), there is no confounding effect of concurrent antidepressant medication. Therefore, any biological changes whether found to be state-independent or state dependent-would entirely reflect the depressive symptom severity while being unaffected by the pharmacological effects of medications.

Analyses were conducted to identify which changes in serum metabolites and pathways were most closely related to changes in depressive symptoms between baseline to week 12 in the acute treatment of MDD with CBT. Herein, we report observed metabolite alterations within a biomarker panel targeting 186 plasma metabolites from 5 distinct metabolite classes (specifically, amino acids, biogenic amines, acylcarnitines, glycerophospholipids, and sphingolipids) available in the Biocrates AbsoluteIDQ^®^ p180 Kit (https://www.biocrates.com/products/research-products/absoluteidq-p180-kit).

## 2. METHOD

### 2.1 Clinical

The PReDICT study was a randomized clinical trial that enrolled 344 adults ages 18–65 years with a primary psychiatric diagnosis of major depressive disorder without psychotic features. The design and clinical results of the study have been published(Dunlop et al. 2012; Dunlop, Kelley, et al. 2017). The Structured Interview for DSM-IV (First et al. 1995) assessed MDD diagnosis, which was confirmed with a psychiatrist’s interview. Patients meeting all eligibility criteria were randomized in a 1:1:1 manner to receive either CBT (delivered in up to 16 one-hour individual sessions), escitalopram, or duloxetine for 12 weeks. One-hundred-fifteen patients were assigned to CBT, of whom 26 had serum samples available for metabolomic analyses at baseline and week 12; these 26 patients are the subjects of the current analysis.

Key inclusion criteria for the trial included no lifetime history of having received treatment for depression (either ≥4 weeks of antidepressant medication at a minimally effective dose or ≥ 4 sessions of an evidence-based psychotherapy), and fluency in either English or Spanish. At screening, patients had to score ≥18 on the HAM-D_17_ (Hamilton 1967) and at the baseline randomization visit had to score ≥ 15. Key exclusion criteria included: a lifetime history of bipolar disorder, psychotic disorder, or dementia; a current significant medical condition that could affect study participation or data interpretation; a diagnosis of obsessive-compulsive disorder, an eating disorder, substance dependence, or dissociative disorder in the 12 months before screening; or substance abuse within the 3 months prior to baseline. The only other psychotropic agents permitted during the trial were sedatives (eszopiclone, zolpidem, zaleplon, melatonin, or diphenhydramine) up to three times per week. The Emory Institutional Review Board and the Grady Hospital Research Oversight Committee approved the study protocol, and all patients provided written informed consent prior to beginning study procedures.

The therapy was delivered in accordance with Beck’s protocol-based CBT(Beck et al. 1979; Beck 1980) and therapists’ fidelity to the protocol was assessed by independent raters at the Beck Institute using the Cognitive Therapy Scale(Anon n.d.). Raters blinded to treatment assignment assessed depression severity using the HAM-D_17_ at baseline, weeks 1-6, 8, 10, and 12. For the individual patient outcomes, the protocol defined remitters as patients who achieved HAM-D_17_ score ≤7 at both week 10 and 12(Dunlop et al. 2012). Consistent with the prior analyses of this dataset(Dunlop, Kelley, et al. 2017), outcomes for non-remitters were using percent change in HAM-D_17_ score from baseline to week 12, as follows:: Non-remitting responder, ≥50% reduction, but not meeting remitter criteria; Partial responder: 30-49% reduction; Treatment failure: <30% reduction.

### 2.2 Laboratory

#### 2.2.1 Metabolomic Profiling using Absolute IDQ p180 Kit

Metabolites were measured with a targeted metabolomics approach using the AbsoluteIDQ^®^ p180 Kit (BIOCRATES Life Science AG, Innsbruck, Austria), with a ultra-performance liquid chromatography (UPLC)/MS/MS system (Acquity UPLC (Waters), TQ-S triple quadrupole MS/MS (Waters)). This procedure provides measurements of up to 186 endogenous metabolites in quantitative mode (amino acids and biogenic amines) and semi-quantitative mode (acylcarnitines, sphingomyelins, phosphatidylcholines and lysophosphatidylcholines across multiple classes). The AbsoluteIDQ^®^ p180 kit has been fully validated according to European Medicine Agency Guidelines on bioanalytical method validation. Additionally, the kit plates include an automated technical validation to assure the validity of the run and provide verification of the actual performance of the applied quantitative procedure including instrumental analysis. The technical validation of each analyzed kit plate was performed using MetIDQ^®^ software based on results obtained and defined acceptance criteria for blank, zero samples, calibration standards and curves, low/medium/high-level QC samples, and measured signal intensity of internal standards over the plate. De-identified samples were analyzed following the manufacturer’s protocol, with metabolomics labs blinded to the clinical data.

#### 2.2.2. Preprocssing of P180 profiles

The raw metabolomic profiles included 182 metabolite measurements of serum samples. Each assay plate included a set of duplicates obtained by combining approximately 10 μl from the first 76 samples in the study (QC pool duplicates) to allow for appropriate inter-plate abundance scaling based specifically on this cohort of samples (n=24 across all plates). Metabolites with >40% of measurements below the lower limit of detection (LOD) were excluded from the analysis (n=160 metabolites passed QC filters). To adjust for the batch effects, a correction factor for each metabolite in a specific plate was obtained by dividing the metabolite’s QC global average by QC average within the plate. Missing values were imputed using each metabolite ‘s LOD/2 value followed by log_2_ transformation to obtain a normal distribution of metabolite levels. The presence of multivariate outlier samples was checked by evaluating the squared Mahalanobis distance of samples. Samples were flagged as “outliers” when their Mahalanobis distances exceeded the critical value corresponding to a Bonferroni-corrected threshold (0.05/n, n: number of samples) of the Chi-square distribution with m degrees of freedom (m=160: number of metabolites).

#### 2.2.3 Data analysis

For statistical analysis we adopted a two-pronged approach. Initially, a multivariate “co-expression network” analysis was employed with CBT treated patients to detect clusters (or modules) of metabolites demonstrating similar patterns of perturbations that correlated with changes in depressive symptom scores based on the HAM-D_17_. Univariate analyses were also performed to detect whether the metabolites within or outside the clusters were individually and significantly correlated to the depressive symptom outcome. The traditional univariate analysis method to metabolomic profiling focuses on the individual metabolites; thus, the interactions among metabolites are largely ignored, even though it is appropriate to assume that metabolites play their roles not in isolation but via interactions with each other. Consequently, metabolite “co-expression” analysis is a powerful, multivariate approach to identify groups of perturbed metabolites belonging to same class or pathways. This approach has the additional benefit of alleviating the multiple testing problem(DiLeo et al. 2011). The workflow of data analysis is presented in **Figure 1**.

**Figure 1:**
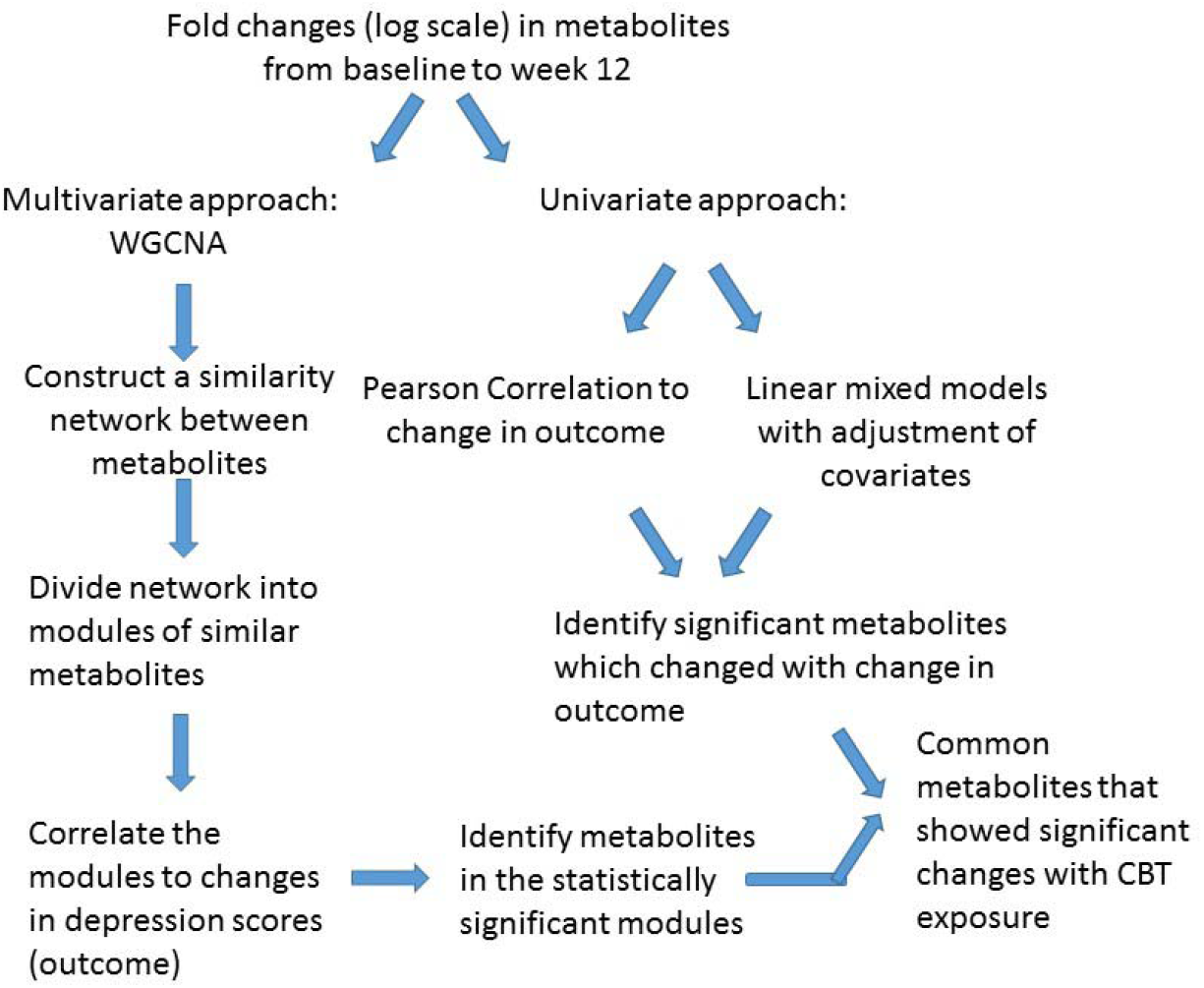
Workflow for data analysis.

##### Univariate Analysis

To define the association between changes in metabolite levels from baseline to week 12 of CBT treatment and the changes in depressive symptom of total HAM-D_17_ scores over that time, linear mixed effects models were fitted to each metabolite change, adjusting for age and gender and with subjects as a random variable. All *p*-values were checked for false discovery rates by Benjamini-Hochberg method(Hochberg and Benjamini 1990). Correlation between metabolite changes and depressive symptoms changes were also assessed by Pearson’s correlation coefficients.

##### Co-expression network analyses

Changes in modules of ‘co-expressed’ metabolites were identified using the R package WGCNA (weighted gene co-expression network analysis)(Langfelder and Horvath 2008). Signed and weighted Pearson’s correlation networks were constructed with the subject-wise changes of baseline to week 12 metabolite concentrations (in the logarithmic scale). First a weighted adjacency matrix was created based on pairwise Pearson’s correlation coefficients between the metabolites. A scale-free topology criterion was used to choose the soft threshold of beta = 18 for the correlations as per the WGCNA protocol. The obtained adjacency matrix was used to calculate the topological overlap measure (TOM) for each pair of metabolite log2 fold changes comparing their adjacencies with all of the other metabolite log2 fold changes. Densely interconnected groups (or modules) of metabolites were identified by hierarchical clustering using 1-TOM as a distance measure through the use of the dynamic hybrid tree cut algorithm with a deep split of 2 and a minimum cluster size of 3. Each module is summarized by the module eigenvector, which is the first principal component of the metabolite changes across all the subjects. Similar clusters were subsequently merged if the correlation coefficient between the clusters’ eigenvectors exceeded 0.75. The association between the resultant modules and the changes in HAM-D_17_ scores was measured by the pairwise Pearson correlation coefficients and presented in a heatmap. In all analyses for this small scale pilot study an uncorrected p-value threshold of 0.10 was used as the significance cutoff.

## 3. RESULTS

### 3.1 Patient Characteristics

Plasma metabolite data were available at baseline and week 12 from 26 patients. The mean number of therapy sessions attended was 14.0 ± 1.5. **Table 1** summarizes the characteristics of the study sample. Twelve (46.2%) of the sample achieved remission and 7 (26.9%) were classified as treatment failures.

**Table 1:** Patient Characteristics

| Measures | Categories | Mean (SD or Percentage) |
| --- | --- | --- |
| No of patients |  | 26 |
| Age<sup>a</sup> |  | 37.4 (10.8) |
| Gender<sup>b</sup> |  |  |
|  | Female | 16 (61.5%) |
|  | Male | 10 (38.5%) |
| Response to Therapy<sup>b</sup> |  |  |
|  | Remitters | 12 (46.2%) |
|  | Responders (Non-remitting) | 3 (11.5%) |
|  | Partial_Responders | 4 (15.4%) |
|  | Treatment Failures | 7 (26.9%) |
| HAM-D<sup>a</sup> |  |  |
|  | Baseline | 18.6 (3.1) |
|  | Week 12 | 8.7 (7.2) |

<sup>a</sup> Mean (SD)
<sup>b</sup> Mean (Percentage)

### 3.2 Detection of metabolite “co-expression” modules

To investigate the functional response of the MDD metabolome during receipt of CBT, we adopted a multivariate approach. Using WGCNA methodology we focused on identifying modules (or clusters) of metabolites that showed a similar pattern of change from baseline to week 12. Thus, each module represented metabolite changes (week12/baseline ratios) in the logarithmic scale. Eight such metabolite modules were identified in which the member metabolites showed statistically significant strong correlations (mean R^2^ ranged between 0.74 and 0.94, all p<0.05) amongst each other in their perturbation patterns, and each module was assigned a unique color. Black, blue, brown, green-yellow, midnight-blue, purple, royal-blue and yellow were the metabolite modules representing 8, 15, 12, 96, 5, 6, 3 and 9 metabolites, respectively. The grey module represented 6 metabolites that could not be assigned to any module. The detected modules were represented by metabolites belonging primarily to the same metabolite class; this may indicate that these metabolites have a functional relationship to each other. Additionally, for each module we identified a ‘hub’ metabolite (also known as a ‘driver’ metabolite) that had the maximum number of connections in the module. The hub metabolites are important and they merit further investigation because they may influence the function of other metabolites, or even may be significant contributors to the trait of interest. The eight modules, their hub metabolites, and their major metabolite classes are presented in **Table 2**. A list of the network metrics, the intra-modular correlations between member metabolites, and the module membership for each metabolite is presented in Supplemental file 1.

**Table 2:** Characteristics of the co-expression modules of metabolites

| Module | Number of member metabolites | Major metabolite classes | Hub metabolite |
| --- | --- | --- | --- |
| Black | 8 | Amino acids | Asn (Asparagine) |
| Blue | 15 | Medium and long chain acylcarnitines | C18:1 (Octadecenoyl-L-carnitine) |
| Brown | 12 | Lysophosphatidylcholines | lysoPC a C18:1 (Lysophosphatidylcholine acyl C18:1) |
| Green-yellow | 96 | Lipids (phosphatidylcholines and sphingomyelins) and acylcarnitines | PC aa C42:4 (Phosphatidylcholine diacyl C42:4) |
| Midnight-blue | 5 | acylcarnitines | C16:2 (Hexadecadienoylcarnitine) |
| Purple | 6 | Short-chain acylcarnitines, α-aminoadipic acid | C3 (Propionyl-L-carnitine) |
| Royalblue | 3 | sphingomyelins | SM C18:1(Sphingomyeline C18:1) |
| Yellow | 9 | Branched-chain amino acids, neurotransmitter-related amino acids | Val (Valine) |

### 3.3 Metabolite modules that were associated with changes in depressive symptoms (HAM-D_17_ scores)

Next, we evaluated the association between the identified metabolite modules and changes in HAM-D_17_ scores from baseline to week 12. Three metabolite modules were found to be significantly associated (R^2^ >0.3, at p<0.1) with changes in the symptom severity scores: a) the purple module containing the short chain acylcarnitines (C3, C4, and C0), α-aminoadipic acid, and the two amino acids, Glutamate and Proline; b) the yellow module containing the BCAAs, Isoleucine and Valine, the BCAA-derived C5-carnitine (Isovalerylcarnitine), the neurotransmitter-related amino acids Tryptophan, Tyrosine, Phenylalanine, Methionine, Methionine-sulphoxide, and the biogenic amine Sarcosine; and c) the green-yellow module containing 96 lipid molecules including the phosphatidylcholines, sphingomyelins, and acylcarnitines. A heatmap showing the correlations between each of the metabolite modules and the changes in HAM-D_17_ scores is presented in **Figure 2**. The correlation coefficients were moderate, ranging between 0.31 and 0.36 for each of the three modules.

**Figure 2:**
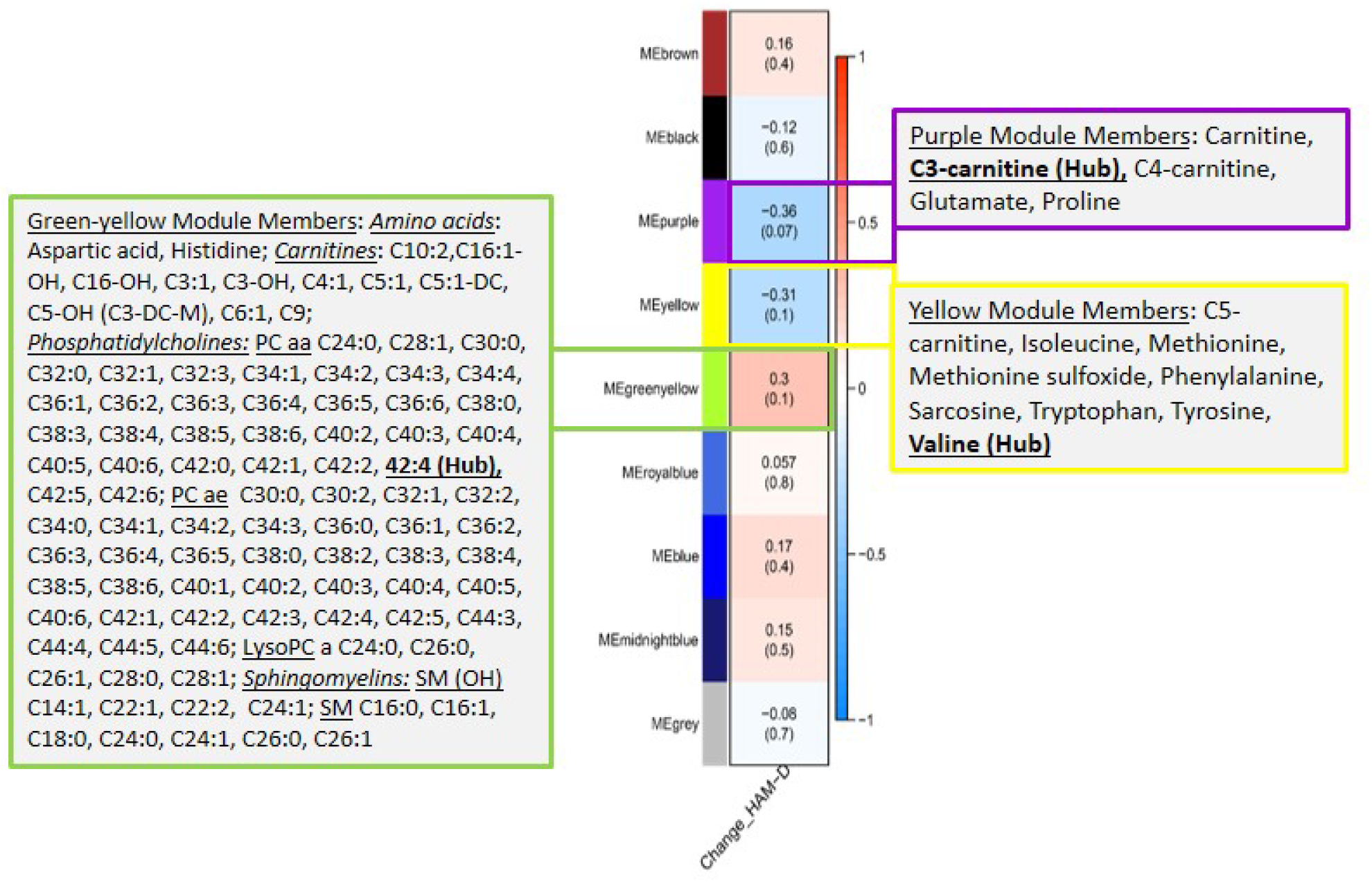
A heatmap showing correlations between the metabolite modules and the changes in symptom severity (HAM-D_17_) scores. Each module represents a number of metabolites that showed strongly correlated perturbation patterns, from baseline to week 12, in response to CBT. The metabolite members of the three modules, purple, yellow and green-yellow, that showed significant association (R^2^ >0.3, at p<0.1) with changes in HAM-D_17_ scores are presented in adjacent boxes.

We examined the correlation of each of the metabolite members of the purple, yellow and green-yellow modules to HAM-D_17_ scores. **Figure 3A** presents a composite plot of the correlations between the changes of each member-metabolite of the purple module to each other and also to HAM-D_17_ changes. Each of the metabolite changes was negatively associated with changes in HAM-D_17_, but they were positively correlated to each other. α-aminoadipic acid showed the strongest correlation to HAM-D_17_ changes (R^2^ = −0.52, p<7E-06). The BCAA-derived C3- (propionyl) and C4- (butyryl) carnitines were strongly correlated to each other (R^2^ = 0.93, p<1.3E-11) as well as to α-aminoadipic acid. C3-carnitine was the hub metabolite in this module.

**Figure 3:**
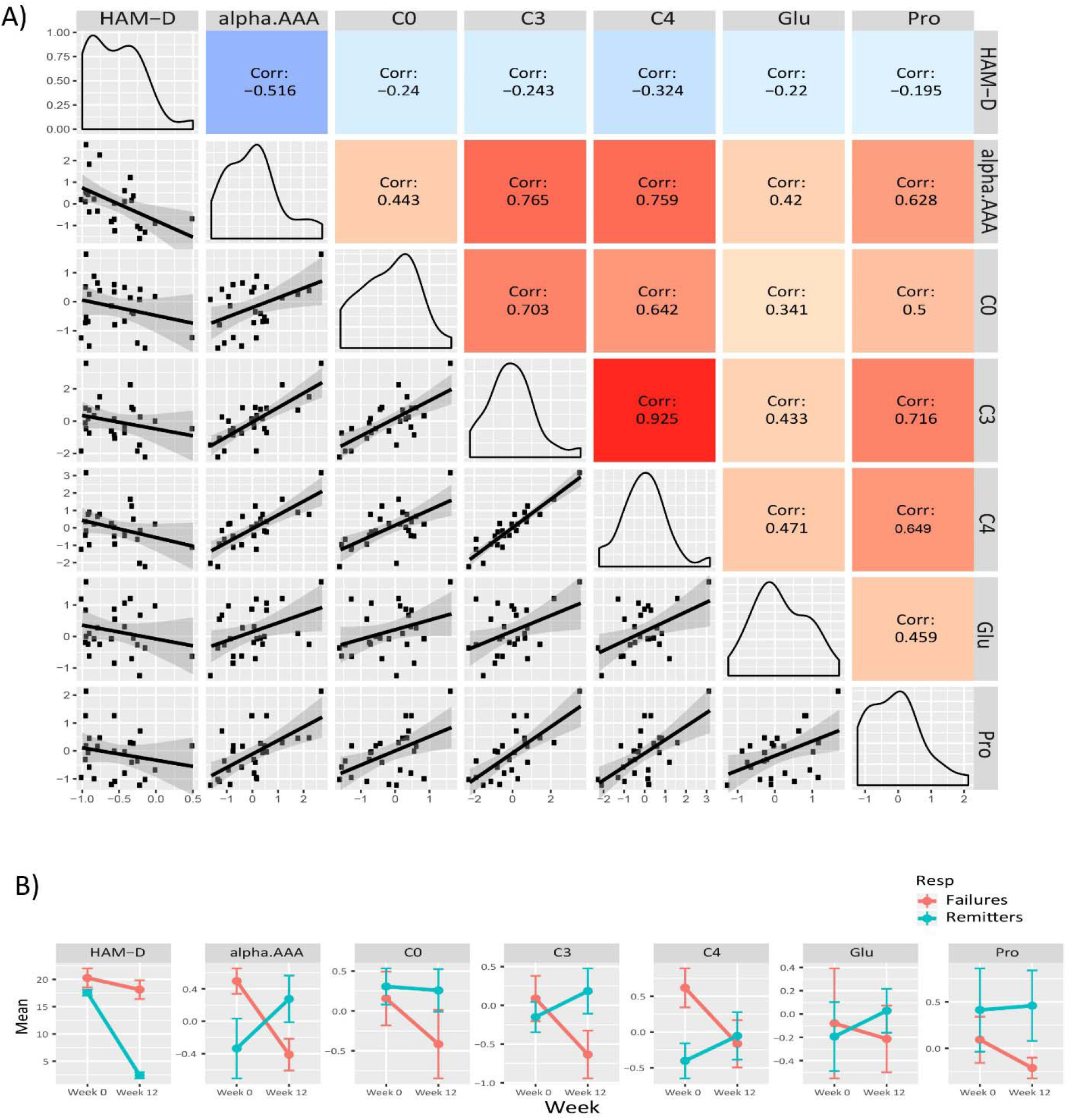
Purple module characteristics. A)The pairwise correlations between the changes in the metabolite members to each other and also to HAM-D_17_ changes are shown in a composite plot. B) The trajectories of each member metabolite with its mean (±SEM) at baseline and week 12 are presented for remitters and treatment-failures.

**Figure 4A** presents a similar composite figure for the yellow module. All members were negatively correlated to HAM-D_17_ changes, with the BCAA valine and the known oxidative stress biomarker methionine sulfoxide perturbations showing the strongest correlations with HAM-D_17_ changes (R^2^ < −0.40, p<0.05). All the yellow module metabolites showed strong positive correlations with each other (mean R^2^ = 0.75, range 0.44 – 0.93, all p< 0.02) in their perturbation patterns. Notable were the strong positive correlations between the branched chain amino acids and methionine, sarcosine, and the neurotransmitter related amino acids (phenylalanine, tryptophan and tyrosine) indicating that they may be functionally related. Valine was the hub metabolite for this module.

**Figure 4:**
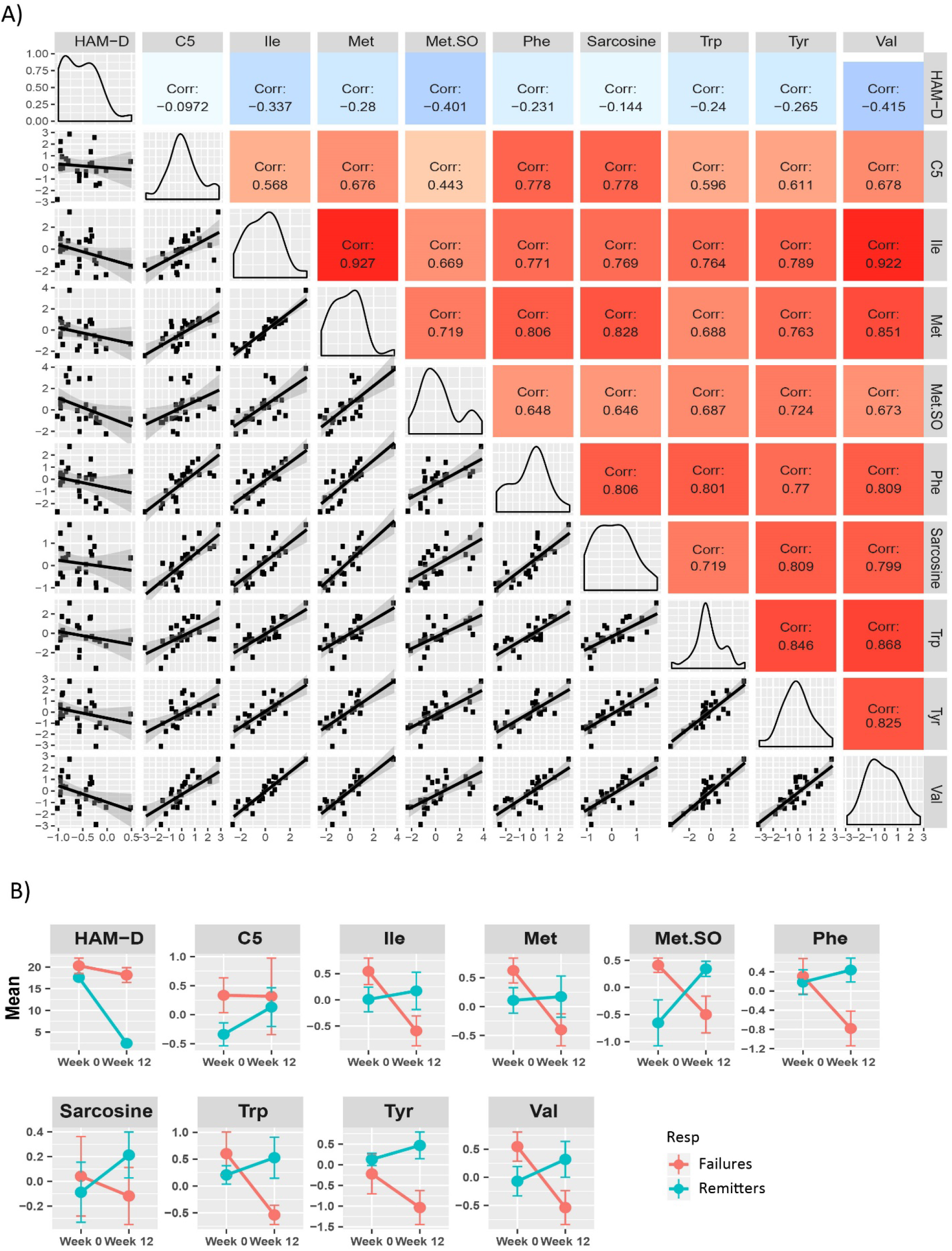
Yellow module characteristics. A)The pairwise correlations between the changes in the metabolite members to each other and also to HAM-D_17_ changes are shown in a composite plot. B) The trajectories of each member metabolite with its mean (±SEM) at baseline and week 12 are presented for remitters and treatment-failures.

The members of the 96 lipids-containing green-yellow module also showed strong correlations amongst each other and also to HAM-D17 changes. Unlike the amino acids and the short chain acylcarnitines from the purple and yellow modules, these lipid molecules’ changes were positively associated with HAM-D_17_ changes and also to each other (Supplemental figures 1A and 1B).

In univariate linear mixed model analysis, the log2 fold changes of the α-aminoadipic acid, the branched chain amino acids, isoleucine and valine, methionine-sulfoxide and several of the phosphatidylcholines, containing long chain fatty acids, from the green-yellow module, were found to be significantly associated (p<0.10) with the changes in HAM-D_17_ scores after adjusting for age and sex. These findings highlight the potential involvements of mitochondrial energy metabolism and lipids in the response of depressed patients receiving CBT. **Table 3** depicts the list of metabolites that were significant in univariate models and their associations with depressive symptom scores (HAM-D_17_).

**Table 3:** Association of metabolite changes to HAM-D_17_ changes from baseline to Week 12 upon exposure to CBT, by univariate models.

| Metabolite | Pearson Correlation |  | Linear mixed model |  | Module-membership |
| --- | --- | --- | --- | --- | --- |
|  | CorrCoeff | Pvalue | CoeffEstimate | Pvalue |  |
| α-aminoadipic acid | -0.5164 | 0.0069 | -0.2 (-0.3, -0.1) | 0.0093 | purple |
| Isoleucine | -0.3372 | 0.0921 | -0.1 (-0.2, 0.0) | 0.0812 | yellow |
| Valine | -0.4147 | 0.0351 | -0.1 (-0.2, -0.0) | 0.0367 | yellow |
| Methionine Sulfoxide | -0.4009 | 0.0424 | -0.1 (-0.2, 0.0) | 0.0512 | yellow |
| Hexenoylcarnitine (C6:1) | 0.3445 | 0.0848 |  |  | green-yellow |
| <sup>a</sup>LPC a C24 | 0.4601 | 0.0180 | 0.1 (0.0, 0.3) | 0.0236 | green-yellow |
| LPC a C26 | 0.4021 | 0.0417 | 0.1 (0.0, 0.2) | 0.0510 | green-yellow |
| LPC a C28 | 0.3666 | 0.0655 | 0.1 (0.0, 0.3) | 0.0730 | green-yellow |
| <sup>b</sup>PC aa C34:1 | 0.4041 | 0.0406 | 0.2 (0.0, 0.3) | 0.0584 | green-yellow |
| PC aa C34:3 | 0.3897 | 0.0491 | 0.1 (0.0, 0.2) | 0.0724 | green-yellow |
| PC aa C36:1 | 0.3501 | 0.0796 |  |  | green-yellow |
| PC aa C36:2 | 0.4238 | 0.0309 | 0.2 (0.0, 0.3) | 0.0431 | green-yellow |
| PC aa C36:3 | 0.4388 | 0.0249 | 0.2 (0.0, 0.3) | 0.0369 | green-yellow |
| PC aa C38:3 | 0.3748 | 0.0592 | 0.1 (0.0, 0.2) | 0.0829 | green-yellow |
| PC aa C38:5 | 0.3479 | 0.0816 |  |  | green-yellow |
| PC aa C40:2 | 0.4232 | 0.0312 | 0.2 (0.0, 0.3) | 0.0419 | green-yellow |
| PC aa C40:3 | 0.3589 | 0.0718 | 0.1 (0.0, 0.3) | 0.0976 | green-yellow |
| PC aa C40:4 | 0.3532 | 0.0767 |  |  | green-yellow |
| PC aa C40:5 | 0.3363 | 0.0930 |  |  | green-yellow |
| PC aa C42:6 | 0.3401 | 0.0891 |  |  | green-yellow |
| <sup>c</sup>PC ae C30:0 | 0.4160 | 0.0346 | 0.2 (0.0, 0.3) | 0.0495 | green-yellow |
| PC ae C34:1 | 0.3875 | 0.0505 | 0.1 (0.0, 0.3) | 0.0702 | green-yellow |
| PC ae C38:2 | 0.5777 | 0.0020 | 0.3 (0.1, 0.4) | 0.0033 | green-yellow |
| PC ae C38:3 | 0.3817 | 0.0543 | 0.1 (0.0, 0.3) | 0.0749 | green-yellow |
| PC ae C40:3 | 0.3604 | 0.0705 | 0.2 (0.0, 0.3) | 0.0952 | green-yellow |
| PC ae C42:1 | 0.4554 | 0.0194 | 0.1 (0.0, 0.3) | 0.0330 | green-yellow |
| PC ae C42:2 | 0.3621 | 0.0691 |  |  | green-yellow |
| PC ae C42:4 | 0.3299 | 0.0998 |  |  | green-yellow |
| Octadecadienylcarnitine (C18:2) | 0.3572 | 0.0732 |  |  | blue |

*Note: According to common lipid nomenclature, <sup>a</sup>LPC stands for lysophosphatidylcholine with a single fatty acid chain, <sup>b</sup>PC aa stands for Phosphatidylcholine diacyl (two fatty acid chains) and <sup>c</sup>PC ae stands for Phosphatidylcholine acyl-alkyl. The lipid species are described as CX:Y where X is the length of the carbon chain C, Y is the number of double bonds; “a” means the acyl chain is attached via an ester bond to the backbone while “e” means the attachment is via an ether bond.*

To maximize our ability to detect the effects of treatment outcomes, we plotted the mean trajectories of the metabolites among the CBT remitters (N = 12) and the treatment failures (N = 7), leaving out the patients with intermediate outcomes, consistent with the approach used in other biomarker studies (Vadodaria et al. 2019; Dunlop, Rajendra, et al. 2017). There were interesting trends among the metabolites in the purple, yellow, and green-yellow modules. In the green-yellow module, consisting of lipids, ~75% of the phosphatidylcholines were higher at baseline in the remitters compared to the treatment failures (Supplemental Figure 2), with some of them being statistically significant at p<0.10 (PC aa- C30:0, C34:1 C36:2, C36:3, PC ae- C36:0, C38:2). In the purple (**Figure 3B**) and yellow (**Figure 4B**) modules, consisting mostly of amino acids including BCAAs and the short-chain acylcarnitines, the remitters mostly had lower baseline levels but showed an upward trend in their metabolite trajectories from baseline to week 12 while the treatment failures all trended to decrease to lower levels post therapy. The limited sample size, however, precluded detection of statistical significance to these differences in trajectories.

## 4. DISCUSSION

Using liquid-chromatography coupled to mass-spectrometry analyses, we examined the biochemical changes that occurred in the plasma of depressed outpatients completing a course of CBT. Changes in several metabolite modules, containing primarily short-chain acylcarnitines and α-aminoadipic acid (purple module) as well as branched-chain and neurotransmitter-related amino acids (yellow module) and lipids (green-yellow module), were significantly associated with changes in depressive symptom severity over the 12-weeks of CBT treatment. The metabolites within each module were highly correlated, and therefore it is likely that the similarity in their perturbations stemmed from their functional relatedness or being members of the same affected pathways.

Changes in individuals’ metabotypes during periods of intense stress, and their return to the original homeostatic levels upon stress resolution (Ghini et al. 2015), support the possibility of identifying metabolomic state markers in MDD. Our analyses found specific amino acids, acylcarnitines, phosphatidylcholines, and sphingomyelins were associated with the depressed state and with changes after CBT treatment. Most notably, concentrations of the BCAAs isoleucine and valine, along with methionine sulfoxide and α-aminoadipic acid, showed strong inverse correlations with change in depression severity. Conversely, many lipid metabolites were directly correlated with changes in depression severity.

Alterations in BCAAs have been linked to altered mitochondrial energy metabolism and have been previously implicated in MDD (Baranyi et al. 2016) and in response to treatment (Kaddurah-Daouk et al. 2013; Kaddurah-Daouk et al. 2011). We have shown that MDD patients have alterations in the phenylalanine, tyrosine, and tryptophan pathways, which are involved in the biosynthesis of the monoamine neurotransmitters ((Bhattacharyya et al. n.d.; Maes et al. 1997; Lucca et al. 1992). Our metabolomic results overlap with the state metabolic markers identified in obesity, type 2 diabetes, and overall worsening metabolic health (Libert et al. 2018; Schooneman et al. 2013). Taken together, the patterns of change observed in our sample implicate bioenergetics as a focus of the pathobiology of the depressed state.

The lipid perturbations, especially those of the phosphatidylcholines (consisting of either diacyl or alkyl-acyl moieties) showed positive correlations to the changes in HAM-D_17_ scores. Phosphatidylcholines are a large class of lipid molecules commonly known as the glycerophospholipids. They have important functions in membrane stability, permeability, and signaling. Phosphatidylcholines have been implicated in MDD in several studies. Recently Knowles *et al*(Knowles et al. 2017) suggested that a subclass of the phosphatidylcholines (the ether-phosphatidylcholines) might have a shared genetic etiology with MDD, and thus might be candidates for improved diagnosis and treatment of depression. Our previous work with ketamine have implicated these lipids along with acylcarnitines in the mechanism of response to ketamine(Rotroff et al. 2016).

These results, suggesting that the metabolites may serve as state markers of depression, received tentative support from our exploratory contrast of the differential trajectories of the changes in metabolites between the patients with the clearest treatment outcomes: remitters versus treatment failures. The BCAAs, their catabolic byproducts, the short chain acylcarnitines, the lysine metabolite α-aminoadipic acid, and the aromatic amino acids (phenylalanine, tyrosine and tryptophan) were present at comparatively higher levels at baseline in the treatment failures compared to the remitters. These findings suggest that metabolic wellbeing may be an important factor contributing to CBT response. Interestingly, there was a general downward trend in the trajectories of these metabolites over the course of treatment among the CBT treatment failures, whereas in the remitters they all exhibited stable or upward trajectories. It is possible that the differences in metabolic trajectories observed in remitters versus treatment failures to CBT indicate a state of “metabolic resilience”(Ghini et al. 2015) in the remitters. The two outcome groups also differed in their metabolomic profiles at baseline, with approximately 75% of the phosphatidylcholines being elevated in the remitters compared to the treatment failures, suggesting that levels of these lipid components may serve as moderators of outcome to CBT. It is also possible that these baseline differences reflect true subtype differences among the MDD patients that might have a role in treatment selection. Larger scale, longitudinal studies will be necessary to test these hypotheses.

The primary limitation to this study is the relatively small number of subjects analyzed. Consequently, testing for the statistical significance of the effects observed was compromised, particularly for the categorical comparisons of differences by treatment outcome group. The study also lacked a healthy control group that could have permitted quantification of how far the observed metabolite concentrations were outside the “normal” ranges. Despite these limtations, the study is important as an original exploration of the metabolomic changes in depression with a proven and well-delivered psychotherapy treatment, in the absence of the powerful metabolomic effects of psychopharmacotherapy.

In summary, this pilot evaluation assessed ~180 metabolites from the Biocrates Absolute p180 kit that clustered into 8 “co-expression modules” based on their propensities to change over 12-weeks of treatment with CBT. The results were largely confirmed by additional univariate analyses of the individual metabolites in the co-expression analyses. Specifically, BCAAs, methionine sulfoxide, α-aminoadipic acid, and multiple phosphatidylcholines were all altered in association with changes in the HAM-D_17_ scores at a level that equaled or exceeded a correlation coefficient of 0.4. Hence, these metabolites may represent markers for the depressed state, and perhaps may act as moderators or mediators for improvement from depression. The results of this study provide an empirically testable set of hypotheses for MDD, namely the utility of these metabolites as potential state markers, in order to understand the mechanistic underpinnings of MDD and their change associated with symptomatic improvement.

## Supporting information

network metrics

the intra-modular correlations between member metabolites

the module membership for each metabolite

## 5. ACKNOWLEDGEMENTS

This work was funded by grant-support to Rima Kaddurah-Daouk through NIH grants R01MH108348, R01AG046171, R01 AG046171 & U01AG061359, RF1AG051550. Sudeepa Bhattacharyya was supported by 5R01MH108348, 5R01AG046171-03S1. Boadie Dunlop was supported by R01MH108348, P50MH077083, R01MH080880, UL1RR025008, M01RR0039 and the Fuqua Family Foundations. Edward Craighead was supported by R01MH080880 and UL1RR025008 (Clinical and Translational Science Award program, NIH) and M01RR0039 (General Clinical Research Center program, NIH). Richard Weinshilboum was supported by R01 GM28157, U19GM61388, U54GM114838 and NSF1624615. Mark Frye was funded by R01MH079261, P20AA017830 and the Mayo Foundation.

## 6. AUTHOR CONTRIBUTIONS

Co-PIs RKD and BWD conceived of the study; SB designed analysis plan; SB and SM performed the data analysis; SB, BWD and AJR wrote the manuscript; SB, BWD, WEC, AJR and RKD assisted in biochemical and clinical interpretations of the results; all authors edited the manuscript.

## 7. CONFLICT OF INTEREST

Boadie Dunlop has received research support from Acadia, Axsome, Intra-Cellular Therapies, Janssen, and Takeda; he has served as a consultant to Myriad Neuroscience and Aptinyx. Richard Weinshilboum is a co-founder and stockholder in OneOme, LLC, a pharmacogenomic clinical decision support company. John Rush has received: consulting fees from Akili, Brain Resource Inc., Compass Inc., Curbstone Consultant LLC., Emmes Corp., Johnson and Johnson (Janssen), Liva-Nova, Mind Linc,, Sunovion, Taj Medical; speaking fees from Liva-Nova; and royalties from Guilford Press and the University of Texas Southwestern Medical Center, Dallas, TX (for the Inventory of Depressive Symptoms and its derivatives). He is also named co-inventor on two patents: U.S. Patent No. 7,795,033: Methods to Predict the Outcome of Treatment with Antidepressant Medication and U.S. Patent No. 7,906,283: Methods to Identify Patients at Risk of Developing Adverse Events During Treatment with Antidepressant Medication. Mark Frye has received grant support from AssureRx Health Inc, Myriad, Pfizer Inc, He has been a consultant (for Mayo) to Janssen Global Services, LLC; Mitsubishi Tanabe Pharma Corp; Myriad Genetics, Inc; Sunovion Pharmaceuticals, Inc; and Teva Pharmaceutical Industries Ltd. He has received continuing medical education, travel, and presentation support from American Physician Institute and CME Outfitters. Dr. Mayberg receives consulting and intellectual property licensing fees from Abbott Labs, Neuromodulation division. Rima Kaddurah-Daouk is inventor on key patents in the field of metabolomics in the study of CNS diseases and holds equity in Metabolon, Inc., a biotechnology company that provides metabolic profiling capabilities. The funders listed above had no role in the design and conduct of the study; collection, management, analysis, and interpretation of the data; preparation, review, or approval of the manuscript; or the decision to submit the manuscript for publication.

