## Supplementary figures and images for "Pilot Study of Metabolomic Clusters as State Markers of Major Depression and Outcomes to CBT Treatment"

### 1B

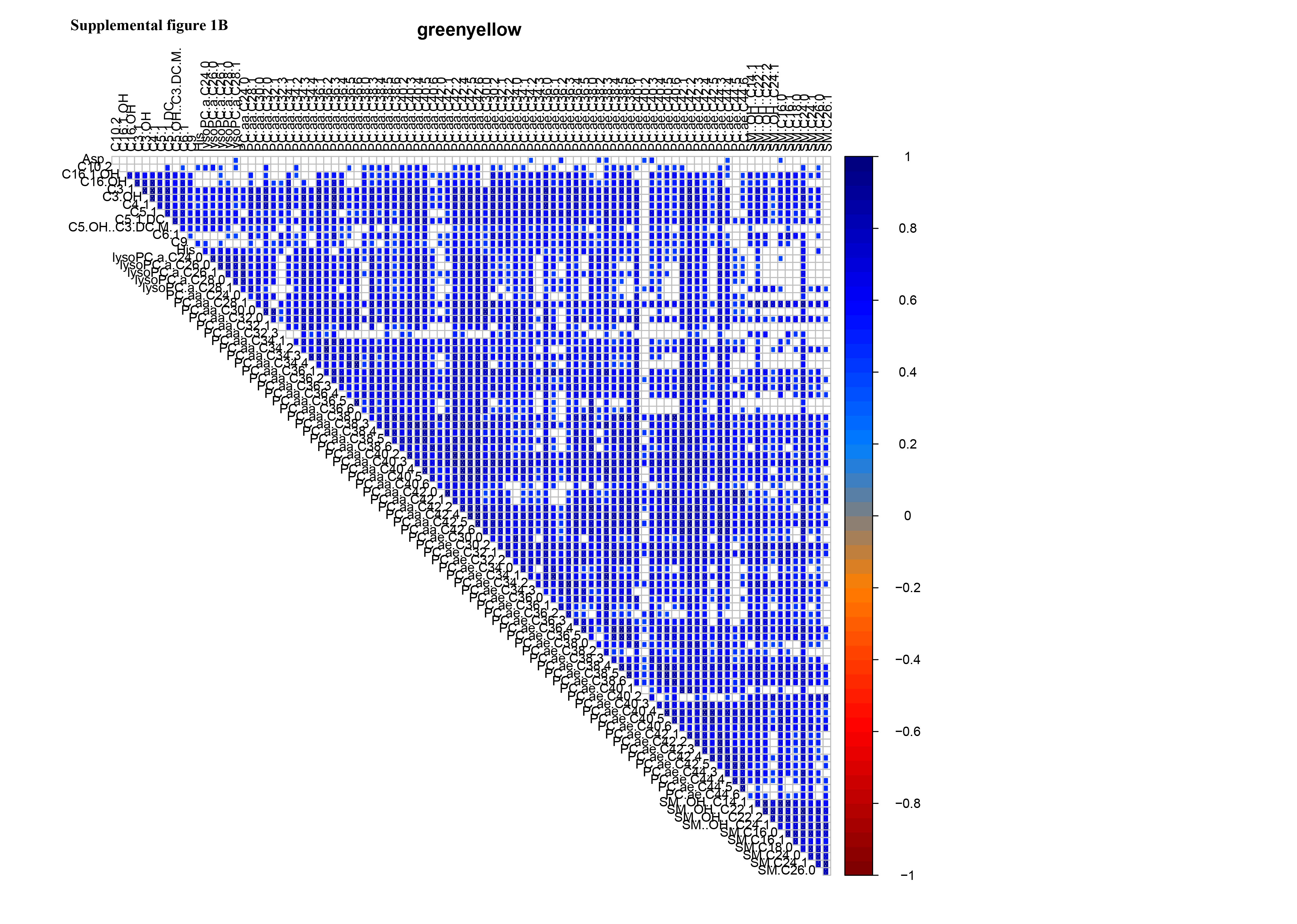

### Supplemental Figure 2

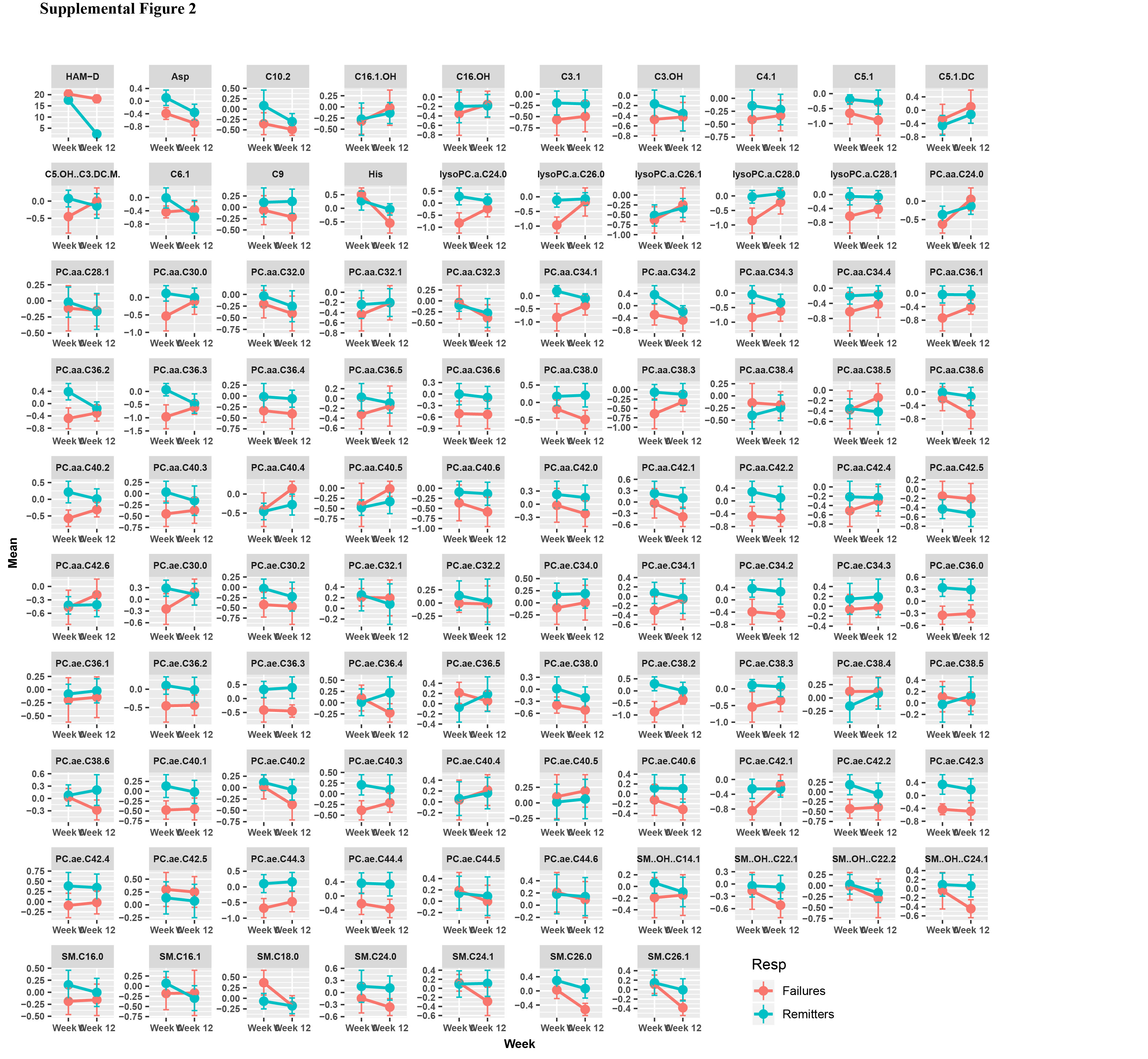

### Supplemental figures 1A

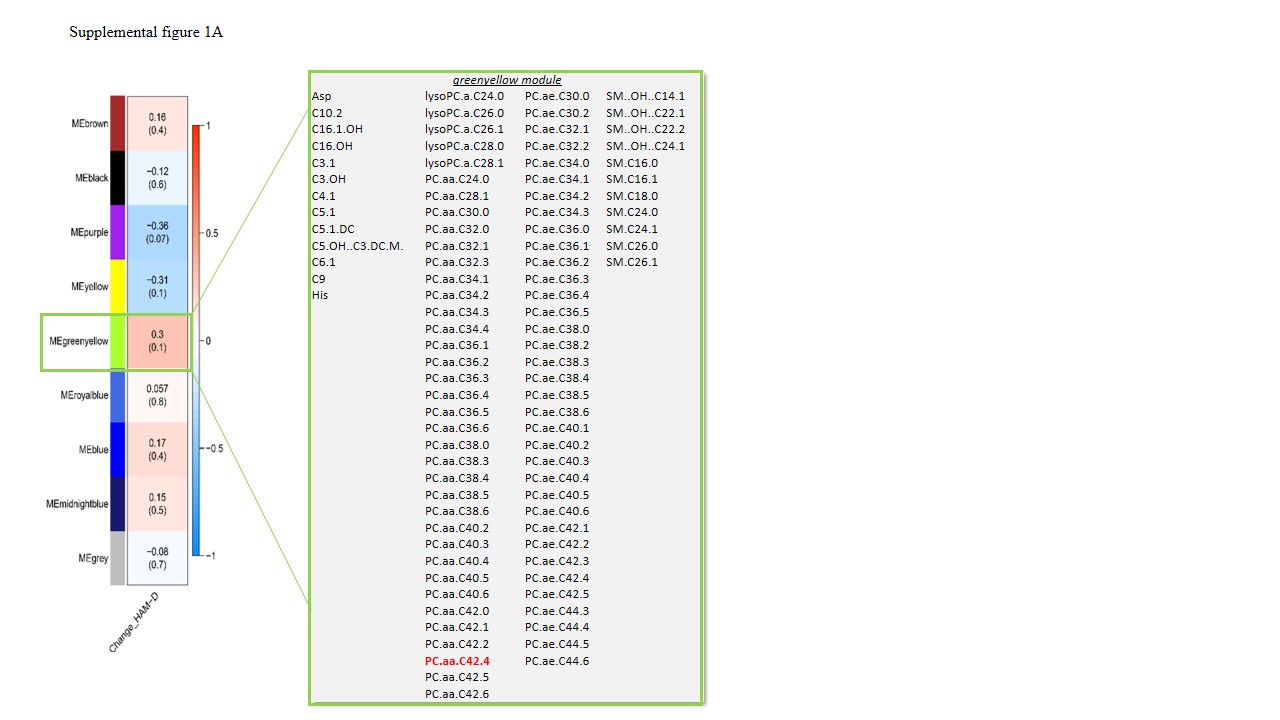
