## Supplemental Figure Legends for "Pilot Study of Metabolomic Clusters as State Markers of Major Depression and Outcomes to CBT Treatment"

Supplemental Figure 1A. A heatmap showing correlations between the metabolite modules and the changes in symptom severity (HAM-D_17)_ scores, highlighting the greenyellow module. The member metabolites of this large module is presented in the adjacent box.

Supplemental Figure 1B. A heatmap showing the pairwise Pearson correlations between the member metabolites of the greenyellow module. The empty cells indicate that the p-values of the correlations for that metabolite-pair was not statistically significant (p>0.1).

Supplemental Figure 2. The trajectories of each member metabolite of the greenyellow module, consisting of mostly lipids, or lipid-related metabolites, with its mean (±SEM) at baseline and week 12 are presented for remitters and treatment-failures.
